# Stochastic models of evidence accumulation in changing environments

**DOI:** 10.1101/019398

**Authors:** Alan Veliz-Cuba, Zachary P. Kilpatrick, Krešimir Josić

## Abstract

Organisms and ecological groups accumulate evidence to make decisions. Classic experiments and theoretical studies have explored this process when the correct choice is fixed during each trial. However, the natural world constantly changes. Using sequential analysis we derive a tractable model of evidence accumulation when the correct option changes in time. Our analysis shows that ideal observers discount prior evidence at a rate determined by the volatility of the environment, and that the dynamics of evidence accumulation is governed by the information gained over an average environmental epoch. A plausible neural implementation of an optimal observer in a changing environment shows that, in contrast to previous models, neural populations representing alternate choices are coupled through excitation.

## Popular Summary

To navigate a constantly changing world, we intuitively use the most recent and pertinent information. For instance, when planning a route between home and work we use recent reports of accidents and weather. We discount older information, as our environment is in constant flux: The clouds threatening rain last night may have dissipated, and an accident reported an hour ago has likely been cleared. How to make decisions in face of uncertainty and impermanence is a question that recurs in fields ranging from economics to ecology and neuroscience. Here, we explore this problem in a general setting where an observer evaluates multiple options based on a series of noisy observations. We assume that the best option is not fixed in time. The optimal strategy is therefore to sequentially update the probability of each alternative, weighting recent evidence more strongly. Our work builds a bridge between statistical decision making in volatile environments and stochastic nonlinear dynamics.

## I. INTRODUCTION

Mammals [1–4], insects [5, 6], and single cells [7] gather evidence to make decisions. However information about the state of the world is typically incomplete and perception is noisy. Therefore, animals make choices based on uncertain evidence. The case of an observer deciding between two alternatives based on a series of noisy measurements has been studied extensively both theoretically and experimentally [2, 8–10]. In this case humans [11], and other mammals [3, 4] can accumulate incoming evidence near optimally to reach a decision.

Stochastic accumulator models provide a plausible neural implementation of decision making between two or more alternatives [12, 13]. These models are analytically tractable [2], and can implement optimal decision strategies [14]. Remarkably, there is also a parallel between these models and experimentally observed neural activity. Recordings in animals during a decision task suggest that neural activity reflects the weight of evidence for one of the choices [3].

However, a key assumption in many models is that the correct choice is fixed in time, *i.e.* decisions are made in a static environment. This assumption may hold in the laboratory, but natural environments are seldom static [15, 16]. Recent experimental evidence suggests that human observers integrate noisy measurements near optimally even when the underlying truth changes. For instance, when observers need to decide between two options and the corresponding reward changes in a history-dependent manner, human behavior approximates that of a Bayes optimal observer [17]. An important feature of evidence accumulation in volatile environments is an increase in learning rate when recent observations do not support a current estimate [18]. Both behavioral and fMRI data show that human subjects employ this strategy when they must predict the position of a stochastically moving target [19]. Experimental work thus suggests that humans adjust evidence valuation to account for environmental variability.

Here we show that optimal stochastic accumulator models can be extended to a changing environment. These extensions are amenable to analysis, and reveal that an optimal observer discounts old information at a rate adapted to the frequency of environmental changes. As a result, the certainty that can be attained about any of the choices is limited. Our approach frames the decision making process in terms of a first passage problem for a doubly stochastic nonlinear model that can be examined using techniques of nonlinear dynamics. This model also suggests a biophysical neural implementation for evidence integrators consisting of neural populations whose activity represents the evidence in favor of a particular choice. Surprisingly, when the environment is not static, these populations are coupled through excitation. This is in contrast to optimal integrators in a static environment [14, 20], and other non-optimal integrators [21, 22] where the populations representing different choices inhibit each other.

## II. OPTIMAL DECISIONS IN A STATIC ENVIRONMENT

We develop our model in a way that parallels the case of a static environment with two possible states. We therefore start with the derivation of the recursive equation for the log likelihood ratio of the two states, and the approximating stochastic differential equation (SDE), when the underlying state is fixed in time.

To make a decision, an optimal observer integrates a stream of measurements to infer the present environmental state. In the static case, this can be done using sequential analysis [1, 9]: An observer makes a stream of independent, noisy measurements, *ξ*_1:*n*_ = (*ξ*_1_*, ξ*_2_,…, *ξ*_*n*_), at equally spaced times, *t*_1:*n*_ = (*t*_1_, *t*_2_,…, *t*_*n*_). The probability of each measurement, *f*_+_(*ξ*_*j*_) := Pr(*ξ*_*j*_*|H*_+_), and *f*_−_(*ξ*_*j*_) := Pr(*ξ*_*j*_*|H*_−_), depend on the environmental state. Combined with the prior probability, Pr(*H*_*±*_), of the states, this gives the ratio of probabilities,

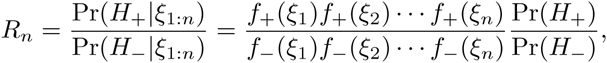

which can also be written recursively [9]:

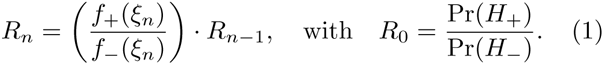

With a fixed number of observations, this ratio can be used to make a choice that minimizes the total error rate [8], or maximizes reward [10]. Eq. (1) gives a recursive relation for the log likelihood ratio, *y*_*n*_ = ln *R*_*n*_,

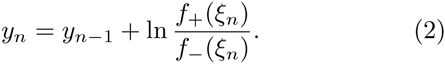

When the time between observations, Δ*t* = *t*_*j*_ – *t*_*j*−1_, is small, we can use the Functional Central Limit Theorem [23] to approximate this stochastic process by the stochastic differential equation (SDE) [2, 24],

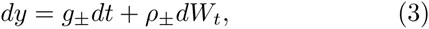

where *W*_*t*_ is a Wiener process, and the constants 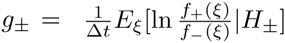 and 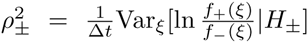 depend on the environmental state. Below we approximate other discrete time process, like Eq. (2), with SDEs. Details of these derivations are provided in the Appendix.

In state *H*_+_ we have 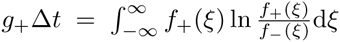. The drift between two observations thus equals the Kullback–Leibler divergence between *f*_+_ and *f*_−_, *i.e.* the strength of the observed evidence from a measurement in favor of *H*_+_. An equivalent interpretation holds for *g*_−_. Hence *g*_+_ and *g*_−_ are the rates at which an optimal observer accumulates information. We will use this observation to interpret the parameters of the model in a changing environment.

**FIG. 1.**
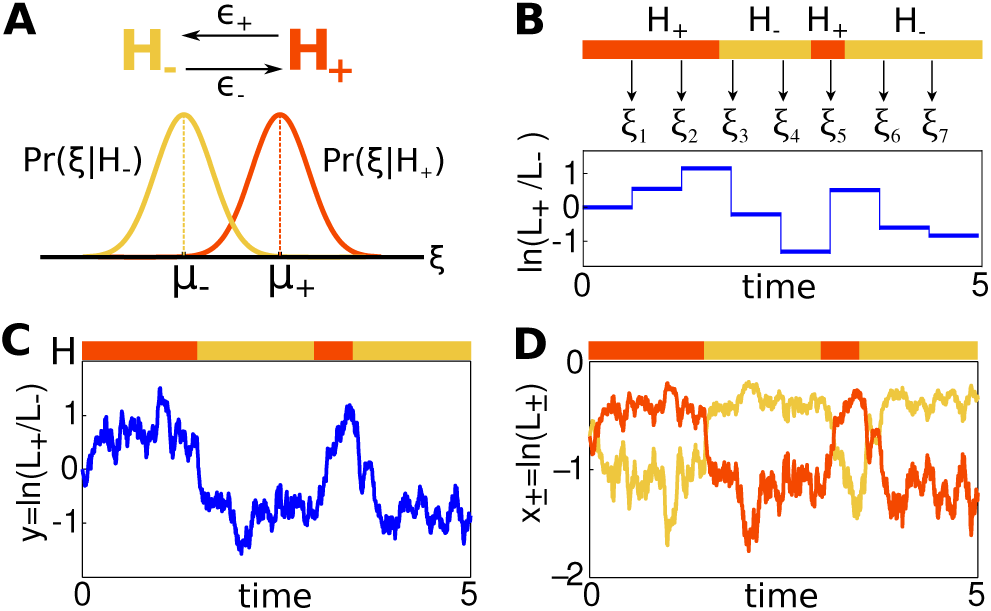
Evidence accumulation in a changing environment. (**A**) The environmental state transitions from state *H*_+_ to *H*_−_ and back with rates *ϵ*_+_, and *ϵ*_−_, respectively. Observations follow state dependent probabilities, *f*_*±*_(*ξ*) = Pr(*ξ H*_±_). (**B**) The distributions of the measurements, *ξ*_*j*_, change with the environmental state. Each individual observation changes the log likelihood ratio, ln(*L*_*n*_,+/*L*_*n*_,-). A single realization is shown. (**C,D**) The evolution of the continuous approximation of the log likelihood ratio, *y*(*t*), (panel **C**) and the log probabilities *x±*(*t*) (panel **D**). At time *t*, evidence favors the environmental state *H*_+_ if *y*(*t*) *>* 0, or, equivalently, if *x*_+_(*t*) > *x*_−_ (*t*).

## III. TWO ALTERNATIVES IN A CHANGING ENVIRONMENT

We use the same assumptions to derive a recursive equation for the log likelihood ratio between to alternatives in a changing environment. The state of the environment, *H*(*t*), is *H*_+_ or *H*_−_, but can now change in time. An observer infers the present state from a sequence of observations, *ξ*_1:*n*_, made at equally spaced times, *t*_1:*n*_, and characterized by probabilities *f*_*±*_(*ξ*_*n*_) := Pr(*ξ*_*n*_ | *H*_*±*_). The state of the environment changes according to a telegraph process [25], and the probability of a change between two observations is *ϵ_±_*Δ*t* := Pr(*H*(*t*_*n*_) = *H*_∓_*|H*(*t*_*n*−1_) = *H*_*±*_). We assume that *ϵ*_+_ and *ϵ*_−_ are known to the observer.

The probabilities, *L*_*n*,±_ = Pr(*H*(*t*_*n*_) = *H*_*±*_*|ξ*_1:*n*_), then satisfy (See Appendix A):

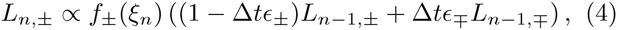

with proportionality constant Pr(*ξ*_1:*n−*1_)*/*Pr(*ξ*_1:*n*_). As in the static case, the ratio of the probabilities of the two environmental states at time *t*_*n*_, can be determined recursively (See Appendix A), and equals

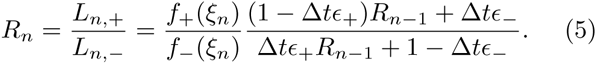

**FIG. 2.**
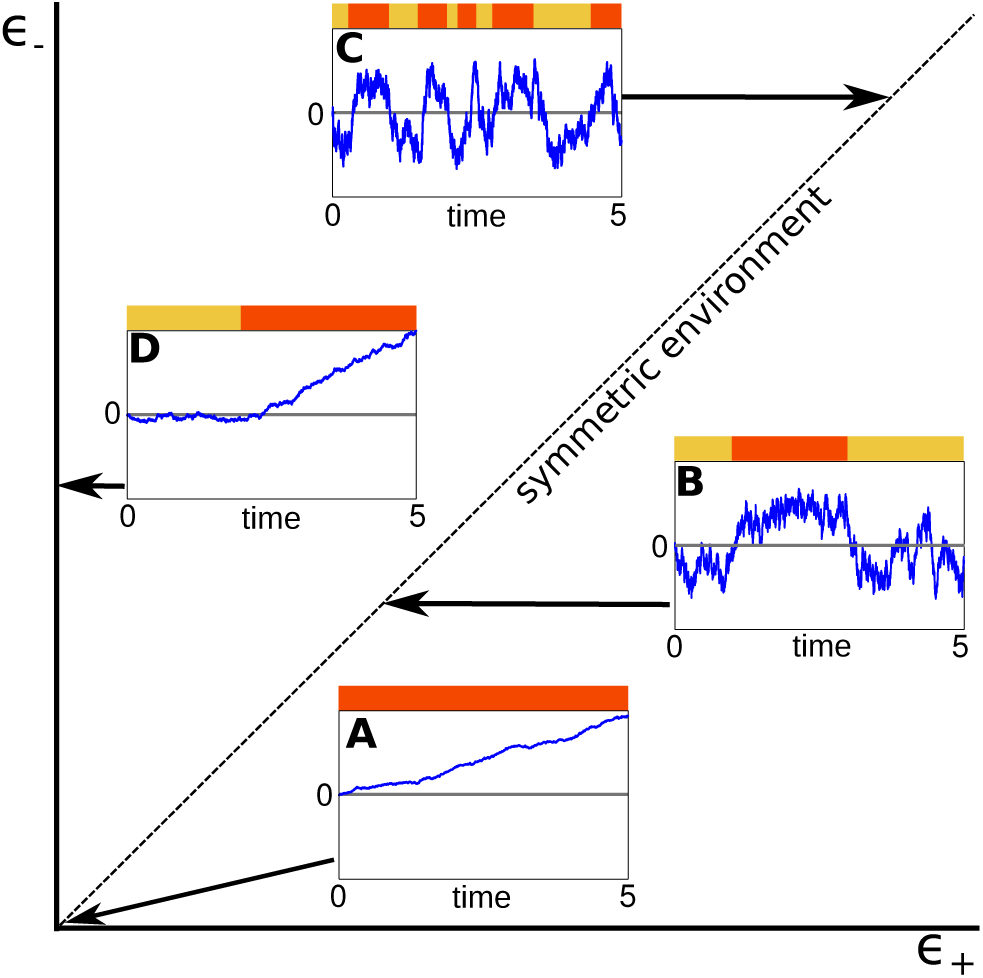
In a dynamic environment, the dynamics of the log likelihood ratio, *y,* depends on the rates of switching between states. (**A**) When *ϵ*_±_ = 0, the environment is static, and the model reduces to the one derived in Section II (**B**) When then environment changes slowly, *|ϵ_±_ | ≪* 1, the log likelihood ratio, *y*, can saturate. (**C**) In a rapidly changing environment, *y* tends not to equilibrate. (**D**) When *ϵ*_+_ = 0 and *ϵ _−_ >* 0, the task becomes a change detection problem.

In this expression, the ratio of probabilities at the time of the previous observations, *R*_*n*−1_, is discounted in a way that depends on the frequency of environmental changes, *ϵ*_±_.

Eq. (5) describes a variety of cases of evidence accumulation studied previously (See Fig. 2): If the environment is fixed (*ϵ*_±_ = 0), we recover Eq. (1). If the environment starts in state *H*_−_, changes to *H*_+_, but cannot change back (*ϵ_−_ >* 0, *ϵ*_+_ = 0), we obtain

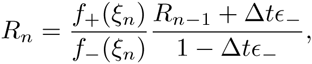

a model used in change point detection [26–28].

We can again approximate the stochastic process describing the evolution of the log likelihood ratio, *y*_*n*_ = ln *R*_*n*_, by an SDE:

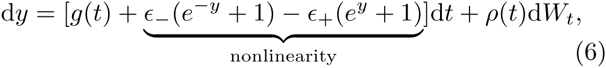

where the drift 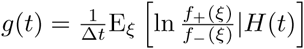, and variance 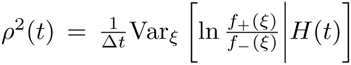 are no longer constant, but depend on the state of the environment at time *t*. We derive Eq. (6) as the continuum limit of the discrete process *y*_*n*_ in Appendix B.

**FIG. 3.**
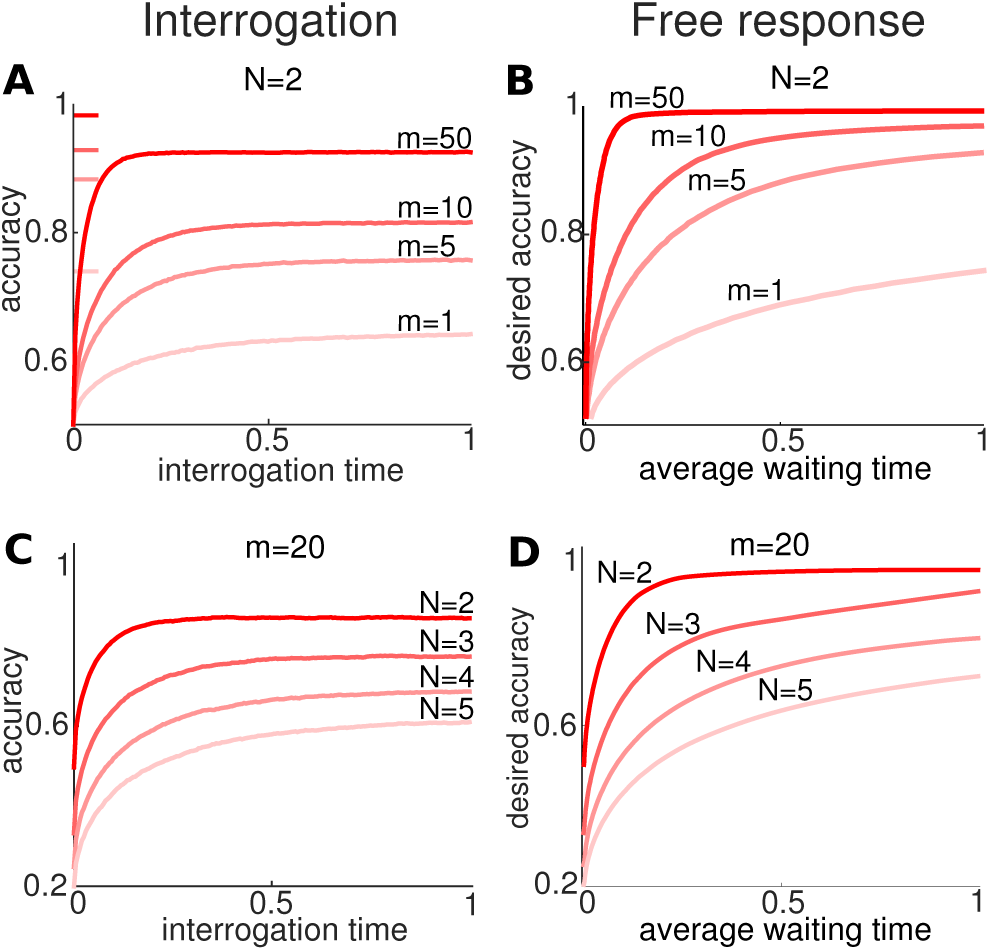
Dependence of the probability of the correct response (accuracy) on normalized information gain, *m,* in a symmetric environment. (**A**) Accuracy in an interrogation protocol increases with *m* and interrogation time, *t,* but saturates. Horizontal bars on left indicate the accuracy when the environment is in a single state for a long time, as in Eq. (10). When the observer responds freely accuracy is similar, but saturates at 1. The increase in accuracy in time is exceedingly slow for low *m*. We fix *ϵ*_*ij*_ ≡ *ϵ* for all *i* ≠ *j*, *g*_*i*_ ≡ *g*, and set *m* = *g*/*ϵ* ≡ 20. (**C**) Accuracy in an interrogation protocol decreases with the number of alternatives *N*, saturating at ever lower levels. (**D**) The free response protocol results in similar behavior, but the accuracy saturates at 1. The increase in accuracy in time is exceedingly slow for higher numbers of alternatives *N*.

The nonlinear term in Eq. (6) does not appear in Eq. (3). It serves to discount older evidence by a factor determined by environmental volatility, *i.e.* the frequency of changes in environmental states. In previous work such discounting was modeled by a linear term [12, 22, 29], however our derivation shows that the resulting Ornstein-Uhlenbeck (OU) process does not model an optimal observer.

### A. Equal switching rates between two states

When *ϵ* := *ϵ*_+_ = *ϵ*_−_, the frequencies of switches between states are equal. Eq. (6) then becomes

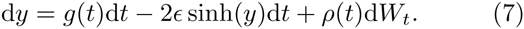

If we rescale time using *τ* = *ϵt*, the rate of switches between environmental states is unity. We obtain

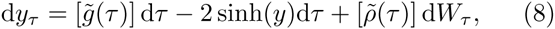

where 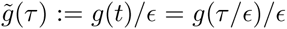 and 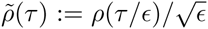. Recall that *g*(*t*) is the rate of evidence accumulation in the present state, and *ϵ*^−1^ is the average time spent in each state. Hence, 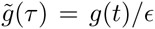 can be interpreted as the *information gained over an average duration of the present environmental state*.

When observations follow Gaussian distributions, *f*_±_ ∼ 𝒩 (±*μ*, *σ*^2^), then *g*(*t*) = *±*2*μ*^2^/*σ*^2^, *ρ* = 2*μ*/*σ*, and

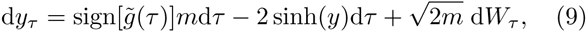

where *m* = 2*μ*^2^/(*σ*^2^*ϵ*). Thus, the behavior of this system is completely determined by the single parameter *m*, the information gain over an average environmental epoch.

The probabilities of a correct response (accuracy) in both interrogation (Fig. 3A) and free response (Fig. 3B) protocols increase with *m*. When an optimal observer is interrogated about the state of the environment at time *τ*, the answer is determined by the sign of the log likelihood ratio, *y*. Since observers discount old evidence at a rate increasing with 1/*m*, decisions are effectively based on a fixed amount of evidence, and accuracy saturates at a value smaller than 1 (Fig. 3A).

If the environment remains in a single state for a long time, the log likelihood ratio, *y*_*τ*_, approaches a stationary distribution,

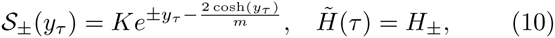

where 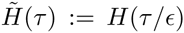 and *K* is a normalization constant. This distribution is concentrated around 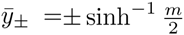, the fixed points of the deterministic counterpart of Eq. (9) obtained by setting *W*_*τ*_ ≡ 0. Since old evidence is continuously discounted, the belief of an optimal observer tends to saturate. In contrast, no stationary distribution exists when *ϵ* = 0, and the environment is static: Aggregating new evidence then always tends to increase an optimal observer’s belief in one of the choices.

Since *𝒮*_±_(*y*) is obtained by assuming that the environment is trapped in a single state for an extended time, 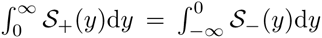 provides an upper bound for accuracy (Fig. 3A). To achieve accuracy *a* in the free response protocol (Fig. 3B), we require 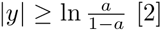 The waiting time for this accuracy steeply increases with *a* and decreases with *m*.

### B. Linear approximation of the SDE

An advantage of Eq. (6) is that it is amenable to standard methods of stochastic analysis. We can find an accurate piecewise linear approximation to Eq. (6), although, for simplicity, we focus on Eq. (9). The piecewise OU process that models an observer that linearly discounts evidence has the form

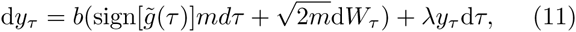

since both drift and diffusion need to be co-scaled. A linear approximation of Eq. (9), with the same equilibria and local stability is obtained by setting 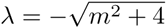 and 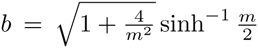. Individual realizations of Eq. (11) and Eq. (9) agree in quickly changing environments (Fig. 4A, *m* = 1), but are less similar in slowly changing environments (Fig. 4B, *m* = 10; see also panel **C**). Thus, as observer performance improves, the nonlinear term in Eq. (9) becomes more important. Note that the corr esponding drift-diffusion model, 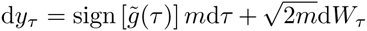, is qualitatively different as it lacks a restorative leak term. This difference becomes more pronounced as *m* increases (Fig. 4C).

**FIG. 4.**
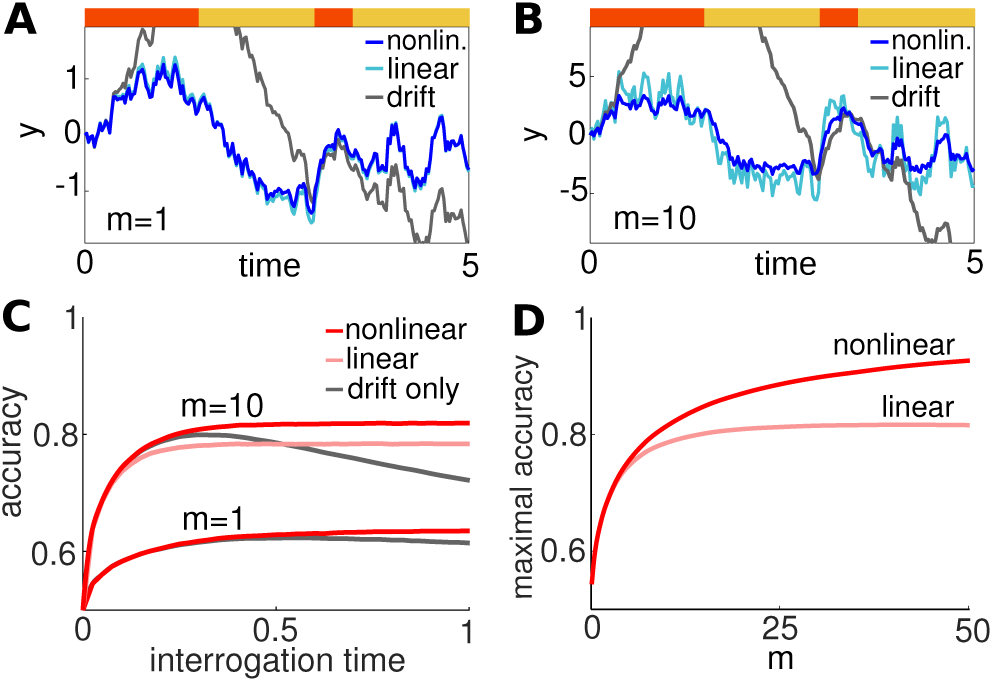
Closest linear approximations of the nonlinear SDE, Eq. (9). (**A**,**B**) Single realizations of the nonlinear Eq. (9), linear approximation Eq. (11), and corresponding drift-diffusion model 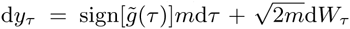, in (**A**) a quickly changing environment (*m* = 1), and (**B**) a slowly changing environment (*m* = 10). We used the same realizations of drift 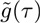, and noise *W*_*τ*_ for all models. (**C**) In the interrogation protocol, accuracy increases faster in the nonlinear Eq. (9) than in the linear approximation Eq. (11). Accuracy eventually decreases in the drift model since all evidence is weighted equally across time. (**D**) In the limit *t* → ∞, accuracy saturates below unity in both the nonlinear model and linear approximation. The linear model employs the discounting strategy of a non-ideal observer, and hence performs worse.

Eq. (11) can be integrated explicitly using standard methods in stochastic calculus [30]. Furthermore, the accuracy of both systems saturate to a value smaller than 1 in the interrogation protocol as the interrogation time increases (Fig. 4C).

The linearized approximation can differ considerably from the full nonlinear model. For instance, in the interrogation protocol the performance of an ideal observer modeled by Eq. (9) increases with interrogation time (Fig. 4D), and accuracy approaches 1 as *m* diverges. In contrast, the accuracy of an observer that discounts evidence linearly limits under 1 as *m* diverges. Indeed, this can be seen by employing the quasi-steady state approximation (fixing sign 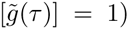, and computing 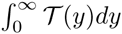 where 𝒯 (*y*) is the steady state distribution of the OU process given in Eq. (11), to obtain

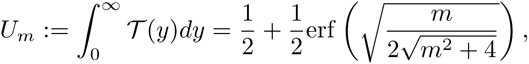

and 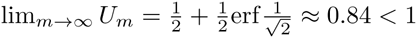

**FIG. 5.**
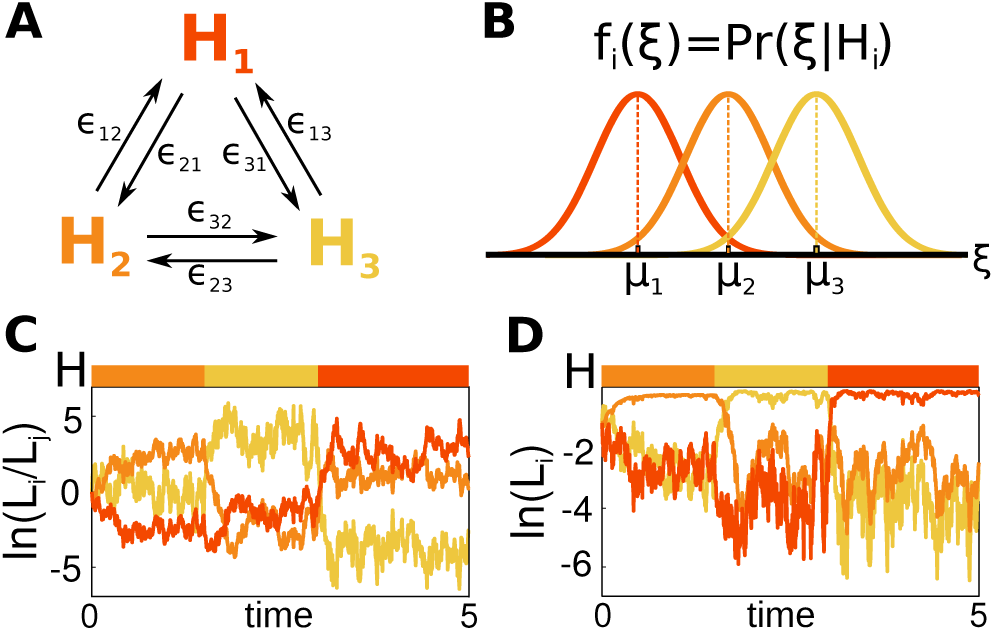
Evidence accumulation with multiple choices in a changing environment. (**A**) The environment switches between *N* states (here *N* = 3). (**B**) Distributions *f*_*i*_(*ξ*) = Pr(*ξ|H*_*i*_) describing the probability of observation *ξ* in environmental state *H*_*i*_(here *N* = 3). (**C,D**) Realization of the log likelihood ratios (panel **C**: ln(*L*_1_/*L*_2_), ln(*L*_2_/*L*_3_), ln(*L*_3_/*L*_1_)), and ln *L*_i_ (panel **D**).

## IV. MULTIPLE ALTERNATIVES IN A CHANGING ENVIRONMENT

We next extend our analysis of evidence accumulation in changing environments to the case of multiple alternatives. With multiple environmental states, *H*_i_ (*i* = 1,…, *N*), the optimal observer computes the present probability of each state from a sequence of measurements, *ξ*_1:*n*_. Measurements have probability *f*_*i*_(*ξ*_*n*_) := Pr(*ξ*_*n*_ *H*_*i*_) dependent on the states *H*_*i*_ [13, 31]. We assume that the state of the environment, *H*(*t*), changes as a memoryless process. A change from state *j* to *i* between two measurements occurs with probability *ϵ*_*ij*_Δ*t* = Pr(*H*(*t*_*n*_) = *H*_*i*_*|H*(*t*_*n*−1_) = *H*_*j*_) for *i ≠ j*, and Pr(*H*(*t*_*n*_) = *H*_*i*_*|H*(*t*_*n*−1_) = *H*_*i*_) = 1 *−* Σ *_j≠i_*Δ*t*ϵ_*ji*_ (Fig 5A).

We again use sequential analysis to obtain the probabilities *L*_*n,i*_ = Pr(*H*(*t*) = *H*_*i*_*|ξ*_1:*n*_) that the environment is in state *H*_*i*_ given observations *ξ*_1:*n*_ The index that maximizes the posterior probability, *î* = argmax_*i*_ *L*_*n*,*i*_, corresponds to the most probable state, given the observations *ξ*_1:*n*_. Following the approach above, we obtain (See Appendix D):

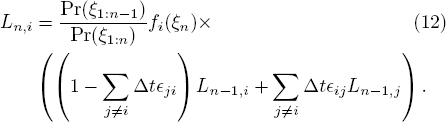

Again after taking logarithms, *x*_*n,i*_ = ln *L*_*n*,*i*_, we can approximate the discrete stochastic process in Eq. (12), with an SDE:

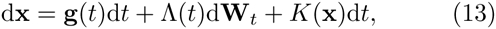

where the drift has components 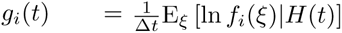, Λ(*t*)Λ(*t*)^*T*^ = Σ(*t*) with entries 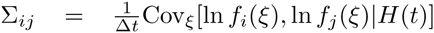, components of **W**_*t*_ are independent Wiener processes, and *K*_*i*_(**x**) = Σ _*j≠i*_ (*ϵ*_*ij*_^e*x*_*j*_^ − ^*x*_*i*_^ − ϵ _*ji*_). The drift *g*_*i*_ is maximized in environmental state *H*_i_.

We can recover the case of two alternatives, *N* = 2, after exchanging the numbers in Eq. (13) with *±* to obtain the approximating SDEs:

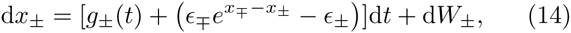

where 〈*W*_*i*_*W*_*j*_〉 = Σ_*ij*_(*t*) *· t* for *i, j ∈ {*+*, −}*. We obtain Eq. (6) by setting *y* = *x*_+_ – *x*_−_. Analogous expressions for the ratios *y*_*ij*_ = ln(*L*_*i*_/*L*_*j*_) are derived in Appendix E. The matrix of these log likelihood ratios quantifies how much more likely one alternative is compared to others (e.g., Fig. 5C) [32].

## V. A CONTINUUM OF STATES IN A CHANGING ENVIRONMENT

Lastly, we consider the case of a continuum of possible environmental states. This provides a tractable model for recent experiments with observers who infer the location of a hidden, intermittently moving target from noisy observations. Evidence suggests that humans update their beliefs quickly and near optimally when observations indicate that the target has moved [19].

Suppose the environmental state, *H*(*t*), intermittently switches between a continuum of possible states, *H*_*θ*_, where *θ* ∈ [*a, b*]. An observer again computes the probabilities of each state from observations, *ξ*_1:*n*_, with distributions *f*_*θ*_(*ξ*_*n*_) := Pr(*ξ*_*n*_ *H*|_*θ*_). The environment switches from state *θ*^′^ to state *θ* between observations with transition probabilities *ϵ_θθ′_* d*θ*Δ*t* := Pr(*H*(*t*_*n*_) = *H*_*θ*_*|H*(*t*_*n*−1_) = *H*_*θ*_ *′*) for *θ ≠ θ*^′^, and 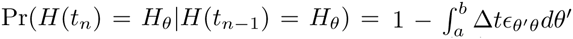 (See Appendix F for details). From Eq. (12) the expression for the probabilities *L*_*n,θ*_ = Pr(*H*(*t*_*n*_) = *H*_*θ*_*|ξ*_1:*n*_) is derived in Appendix F, yielding:

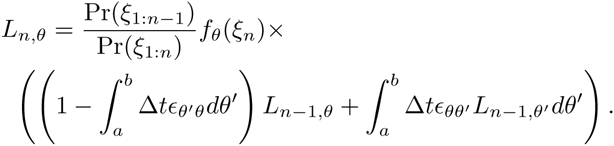

**FIG. 6.**
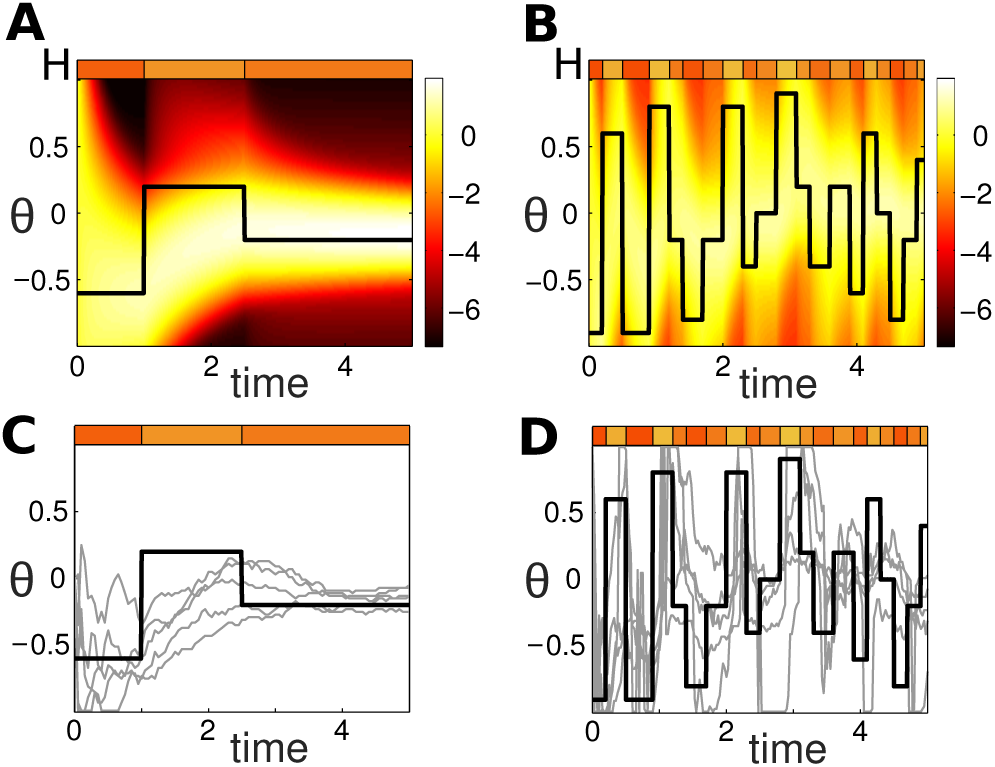
Evidence accumulation with a continuum of choices. The observer infers the state of the environment, *H*_*θ*_, where *θ* ∈ [*−*1, 1], and the state changes at discrete points in time. (**A**) In slowly changing environments, the distribution of the log probabilities, *x*_*θ*_, can nearly equilibrate between switches (Solid line represents the true state of the environment at time *t*. For clarity, we show results of simulations without noise). (**B**) In quickly changing environments, the distribution does not have time to equilibriate between switches. (**C**) In slowly changing environments, the most probable state of the environment, 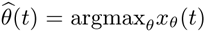 (thin lines), fluctuates around the true value (thick line). (**D**) In quickly changing environments, 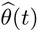 fluctuates more widely, as it is in a transient state much of the time.

We again approximate the logarithms of the probabilities, ln *L*_*n,θ*_, by a temporally continuous process,

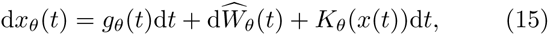

where, *x* = (*x*_θ_)_θ∈[*a*, *b*]_, 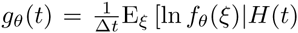, Ŵ_θ_ is a spatiotemporal noise term with mean zero and covariance function given by

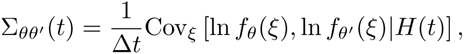

and 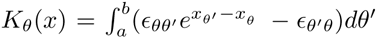 is an interaction term describing the discounting process.

The drift *g*_*θ*_(*t*) is maximal when *θ* agrees with the present environmental state. The most likely state, given observations up to time *t*, is 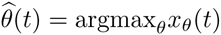.

In slowly changing environments, the log probability *x*_*θ*_(*t*) *nearly* equilibrates to a distribution with a welldefined peak between environmental switches (Fig. 6A). This does not occur in quickly changing environments (Fig. 6B). However, each logarithm, *x*_*θ*_(*t*) approaches a stationary distribution if the environmental state remains fixed for a long time. The term *K*_*θ*_(*x*) in Eq. (15) causes rapid departure from this quasi-stationary density when the environment changes, a mechanism proposed in [19].

Even when the environment is stationary for a long time, noise in the observations stochastically perturbs the log probabilities, *x*_*θ*_(*t*), over the environmental states. This leads to fluctuations in the estimate 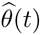 of the most probable alternative (Fig. 6C, D). Thus, as opposed to the case of a discrete space of *N* alternatives, the observer’s estimate of the most probable choice will change continuously, fluctuating about the continuum of possible alternatives. Unless changes are too rapid, the peak of the log probability distribution, 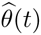, fluctuates around the true environmental state, and tracks abrupt changes in *H*_*θ*_(*t*). This is in line with recent observations in human behavioral data [19].

**FIG. 7.**
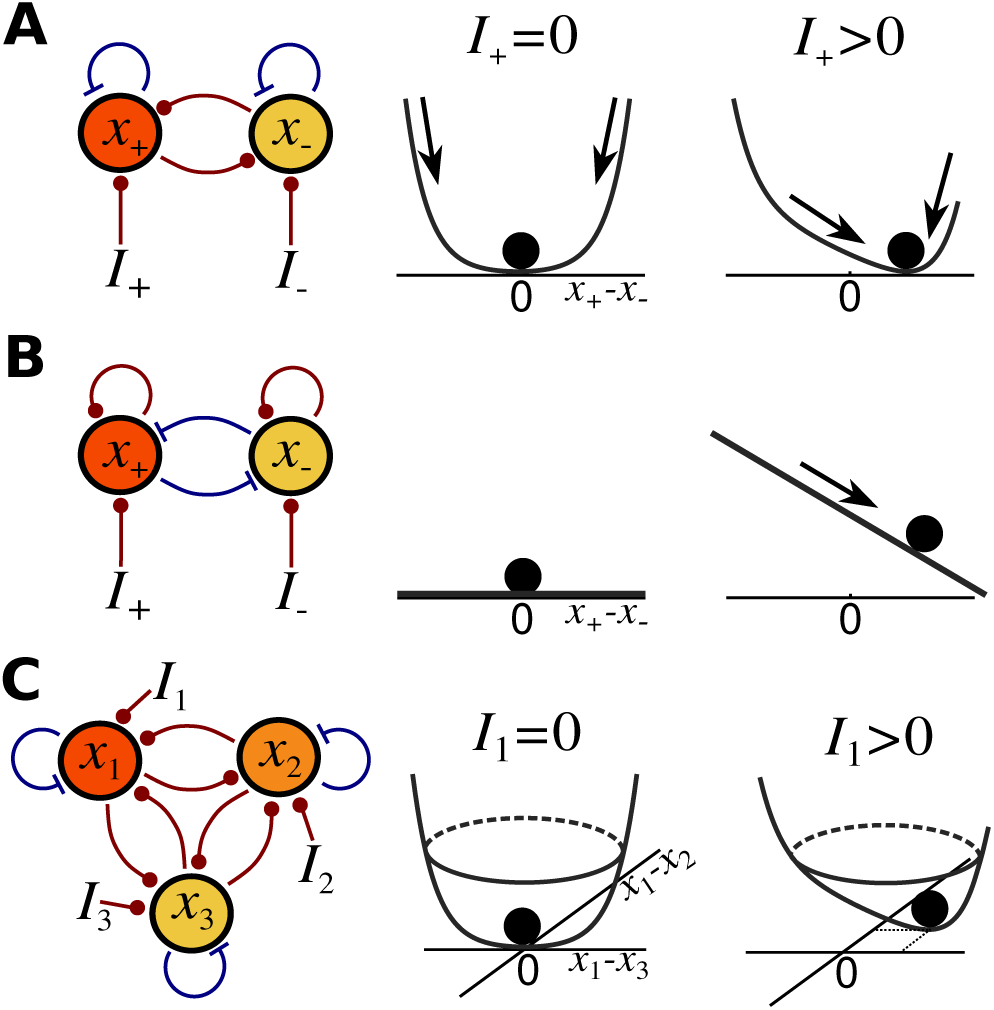
Neural population models of evidence accumulation. (**A**) Two populations *u*_*±*_ receive a fluctuating stimulus with mean *I*_*±*_; they are mutually coupled by excitation (circles) and locally coupled by inhibition (flat ends). When *I*_+_ *>* 0, the fixed point of the system has coordinates satisfying *x*_+_ *> x_−_* as shown in the plots of the associated potentials. (**B**) Taking *ϵ_±_ →* 0 in Eq. (16) generates a mutually inhibitory network that perfectly integrates inputs *I*_*±*_ and has a flat potential function. (**C**) With *N* = 3 alternatives, three populations coupled by mutual excitation can still optimally integrate the inputs *I*_1,2,3_, rapidly switching between the fixed point of the system in response to environmental changes.

## VI. A NEURAL IMPLEMENTATION OF AN OPTIMAL OBSERVER

Previous neural models of decision making typically relied on mutually inhibitory neural networks [12, 20, 33], with each population representing one alternative. In contrast, inference in dynamic environments with two states, *H*_+_ and *H*_−_, can be optimally performed by mutually excitatory neural populations with activities (firing rates) *r*_+_ and *r*_−_,

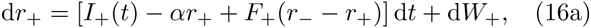

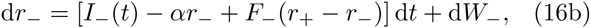

where the transfer functions are *F*_*±*_(*x*) = *−αx/*2+*ϵ_∓_*e^*x*^-ϵ_*±*_, the mean input 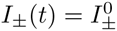 when *H*(*t*) = *H*_*±*_ and vanishes otherwise, *W*_*±*_ are Wiener processes representing the variability in the input signal with covariance defined as in Eq. (14) (See Appendix D). Thus, *I*_*±*_(*t*)d*t* + d*W*_*±*_.represents the total input to population *r*_*±*_. When *α >* 0 and sufficiently small, population activities are modulated by self-inhibition, and mutual excitation (Fig. 7A). Note, the parameter *α* determines the leak in the activity of each individual population, which depends on both the time constants and recurrent architecture of the local network [33]. Taking *y* = *r*_+_ *− r_−_* reduces Eq. (16) to the SDE for the log likelihood ratio, Eq. (6). In the limit of a stationary environment, *ϵ_±_ →* 0, we obtain a linear integrator d*r*_*±*_ = [*I*_*±*_d*t* + d*W*_*±*_] *− α*(*r*_+_ + *r*_−_)d*t/*2, as in previous studies [2, 20].

To show that the populations mutually excite each other, we set *W*_+_ = *W*_−_ = 0, and linearize Eqs. (16). When the environment has not changed for a long time, Eq. (16) approaches a fixed point 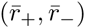 with

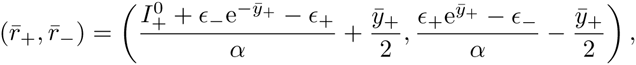

when 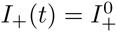 and *I*_−_(*t*) = 0 and

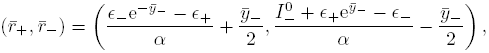

when *I*_+_(*t*) = 0 and 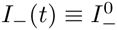, where

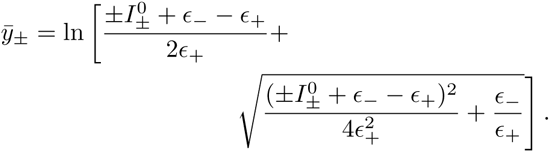

Note that by increasing (decreasing) *α*, the fixed points 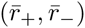 move closer to (farther from) the origin (0, 0). To determine the sign of the coupling near these fixed points, note that the Jacobian matrix of (*F*_+_*, F_−_*) has the form:

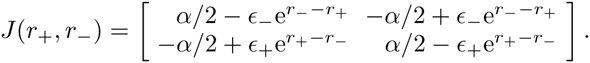

For *ϵ_±_ >* 0, taking *α <* 2 min 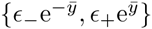 will guarantee that the sign of the Jacobian matrix is 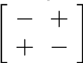 on a region that contains the fixed point. This corresponds to a neural network with self-inhibition and mutual excitation as illustrated in Fig. 7A.

The model given by Eq. (16) is matched to the timescale of the environment determined by *ϵ_±_.* Solutions approach stationary distributions if input is constant. Due to mutual excitation, they are very sensitive to changes in inputs, a feature absent in previous connectionist models [34]. Even when *E* is small, Eq. (16) has a single attracting state determined by the mean inputs 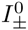. We illustrate the response of the model to inputs using potentials (Fig. 7A). In contrast to the single attractor of Eq. (16), mutually inhibitory models can possess a neutrally stable line attractor that integrates inputs (*ϵ_±_ ≡* 0, Fig. 7B) [35].

We can extend our results for the *N* = 2 case by deriving a neural population model of decision making in changing environments. In [13], the reliability of motion information was assumed to vary during a trial, and the optimal model encoded the posterior probability distribution over the possible stimulus space. Here, we assume the true hypothesis, *H*(*t*), changes in time. For an arbitrary number of possible states, {*H*_1_,…, *H*_*N*_}, decisions can be performed optimally by neural populations *x*_1_,…, *x*_*N*_ coupled by mutual excitation

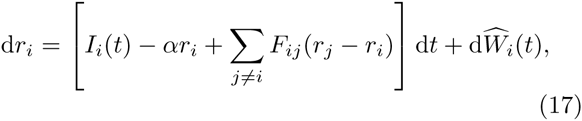

where the mean input is 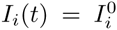 when *H*(*t*) = *H*_*i*_ and 0 otherwise and (d Ŵ_1_(*t*)*,…,* dŴ_*N*_ (*t*))^*T*^ = Λ(*t*)d**W**_*t*_ describes input noise with Λ(*t*) defined as in Eq. (13). Population firing rates are again determined by inhibition within each population and excitation between populations as described by the arguments of the firing rate function

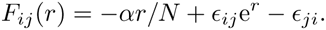

In this case coupling between populations is again excitatory (Fig. 7C).

Note that, as in the case of *N* = 2 alternatives, taking the limit of Eq. (17) as *ϵ*_*ij*_ → 0, we obtain linear integrators [20]

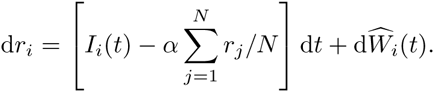

## VII. DISCUSSION

We have derived a nonlinear stochastic model of optimal evidence accumulation in changing environments. Importantly, the resulting SDE is not an OU process, as suggested by previous heuristic models [12, 22, 24]. Rather, an exponential nonlinearity allows for optimal discounting of old evidence, and rapid adjustment of decision variables following environmental changes. As a result, the certainty of an optimal observer tends to saturate, even if the environment happens to be stuck in a single state for an extended time.

We have made several assumptions about the model to simplify these initial derivations. Our ideal observer is assumed to be aware both of the uncertainty of their own measurements and about the frequency with which the environment changes. A more realistic model would require that a naive observer learn the underlying volatility of the environment. Efforts to model the case of initially unknown transition rates produced hierarchical models that identify the location of change-points [36]. However, this approach quickly grows in computational complexity, since the probability of change points is determined by accounting for all possible transition histories [27]. We also assumed that changes in the environment follow a memoryless process. In more general cases, we would not be able to obtain a recursive equation for the probability of a state. An ideal observer would have to use all previous observations at each step, rather than integrating the present observation with the posterior probability obtained with the previous observation. This process cannot be approximated by an SDE.

Sequential sampling in dynamic environments with two states has been studied previously in special cases, such as adapting spiking models, capable of responding to environmental changes [37]. Likelihood update procedures have also been proposed for multiple alternative tasks in the limit *ϵ*_*ij*_ *→* 0 [32, 38]. Furthermore, Eq. (12) for the case *N* = 2 was derived in [39], but its dynamics were not analyzed. One important conclusion of our work is that *m* = *g/ϵ*, the information gain over the characteristic environmental timescale, is the key parameter determining the model’s dynamics and accuracy. It is easy to show that equivalent parameters govern the dynamics of likelihoods of multiple choices. This allows for a straightforward approximation of the nonlinear model by a linear SDE, which can be analyzed fully.

Models of evidence accumulation are not commonly discussed in the physics literature, but are of interest in disciplines ranging from neuroscience and robotics to psychology and economics. They can help us understand how decisions are made in cells, animals, ecological groups, and social networks. We presented a principled derivation of a series of nonlinear stochastic models amenable to stochastic analysis, and have used quasistatic approximations, first passage techniques, and dimensional analysis to examine their dynamics. Thus we have built a bridge between classic models in signal detection theory and nonlinear stochastic processes. Continuous stochastic models have been very useful in interpreting human decision making in static environments [2, 3]. Dynamic environments offer a promising future direction for theory and experiments to probe the biophysical mechanisms that underlie decisions.

## Acknowledgements

Funding was provided by NSF-DMS-1311755 (ZPK); NSF/NIGMS-R01GM104974 (AV-C and KJ); and NSF-DMS-1122094 (KJ).

## APPENDIX

In this appendix, we present the derivations for the probability update formulas and their approximations discussed in the main text. We begin by deriving the update expression for the probability ratio, *R*_*n*_, in the case of two alternatives in a changing environment. The result is a nonlinear recursive equation. Subsequently, we show how to approximate the log likelihood ratio, *y*_*n*_ = ln *R*_*n*_, using a SDE. To make the approximation precise, it is key to view the discrete equation for *y*_*n*_ as a family of equations parameterized by the time interval, Δ*t,* over which each observation, *ξ*_*n*_, is made [2]. Furthermore, we extend our derivations to multiple (*N >* 2) alternatives, and show that the log probability updates can be approximated by a nonlinear system of SDEs in the continuum limit. With the appropriate scaling of the probabilities, *f*_*i*_(*ξ*) = Pr(*ξ*|*H*_*i*_), we can make precise the correspondence between the discrete and continuum models of posterior probability evolution. Lastly, we present a derivation for the stochastic integro–differential equation that represents the log probability for a continuum of possible environmental states, *θ* ∈ [*a, b*].

Note that throughout the appendix, we use notation involving a subscript Δ*t*. This helps us define a family of stochastic processes indexed by the spacing between observations Δ*t* = *t*_*n*_ − *t*_*n*−1_. For instance, *f*_Δ*t*,±_(*ξ*) represents the probability of an observation, *ξ,* in environmental state *H*_±_ (or, in the language of statistics, when hypothesis *H*_±_ holds). This probability changes with the timestep Δ*t*. This approach allows us to properly take the continuum limit Δ*t →* 0. However, for simplicity we refrain from using this notation in the main text. Rather, we treat the limiting SDEs as approximations of discrete update processes. Also, we slightly abuse notation and write *f*_*i*_(*ξ*) = Pr(*ξ*|*H*_*i*_), even when *ξ* is a continuous random variable.

### Appendix A: Likelihood ratio for two alternatives

We begin by deriving the recursive update equation for the probabilities *L*_*n,±*_ := Pr(*H*(*t*_*n*_) = *H*_±_|*ξ*_1:*n*_) associated with each alternative *H*_*±*_, where each observation (measurement), *ξ*_*i*_, is made at time *t*_*i*_. This is the probability that alternative *H*_*±*_ is true at time *t*_*n*_, given that the series of observations *ξ*_1:*n*_ has been made. Importantly, the underlying truth changes stochastically, and in a memoryless way, with transition probabilities given by *ϵ*_Δ*t,±*_ := Pr(*H*(*t*_*n*_) = *H*_∓_*|H*(*t*_*n*−1_) = *H*_*±*_), so that Pr(*H*(*t*_*n*_) = *H*_*±*_ *H*(*t*_*n*−1_) = *H*_*±*_) = 1 *−ϵ*_Δ*t,±*_. We begin by examining the probability *L*_*n*,+_ associated with the alternative *H*_+_. Using Bayes’ rule and the law of total probability we can relate the current probability, *L*_*n*,+_, to the conditional probabilities at the time of the previous observation, *t*_*n−*1_:

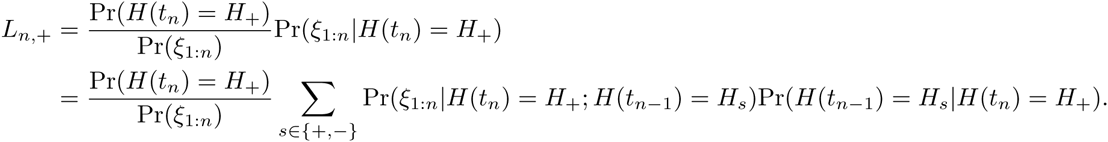

To derive a recursive equation, with probabilities that are not conditioned on the state at *t*_*n*_, we first use Bayes’ rule again to write

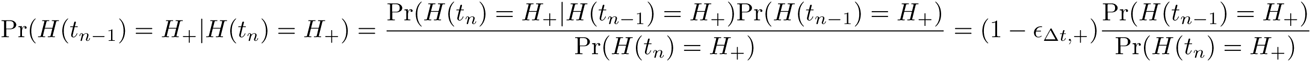

and

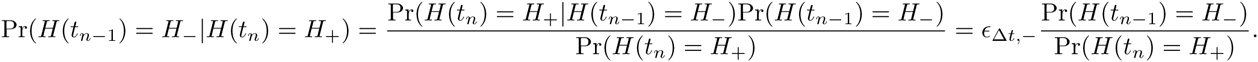

Plugging these formulas into our expression for *L*_*n,*+_, we can then write

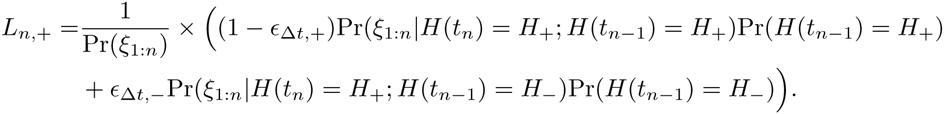

The observation *ξ*_*n*_ is independent from the sequence of observations *ξ*_1:*n−*1_ when conditioned on the states *H*(*t*_*n*_) = *H*_+_ and *H*(*t*_*n−*1_) = *H*_*±*_, respectively. Thus, we obtain

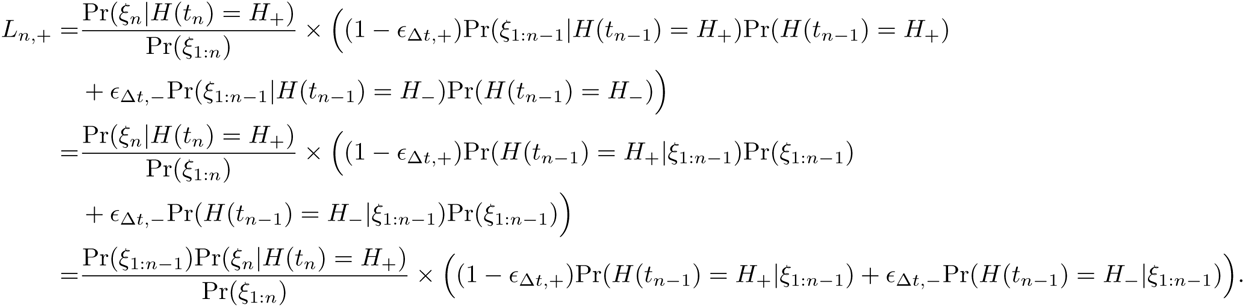

Thus, by using our definition of the probabilities *L*_*n,±*_, we can write an update equation for *L*_*n,*+_ in terms of the probabilities *L*_*n−*1,*±*_ at the previous time, *t*_*n−*1_,

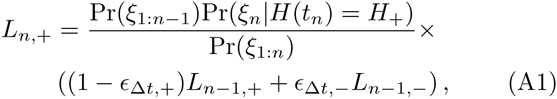

where *L*_0,+_ = Pr(*H*_+_, *t*_0_).

Similarly we obtain an update equation for the probability *L*_*n*,−_ of the alternative *H*_−_ at time *t*_*n*_:

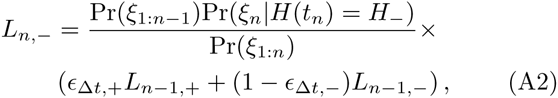

where *L*_0,-_ = Pr(*H*_−_, *t*_0_).

From Eqs. (A1) and (A2), the ratio *R*_*n*_ = *L*_*n*,+_/*L*_*n*,-_ is readily seen to satisfy the recursive equation

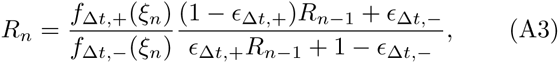

where *f*_Δ*t,±*_(*ξ*_*n*_) = Pr(*ξ*_*n*_ *H*(*t*_*n*_) = *H*_*±*_) is the distribution for each choice parameterized by the timestep Δ*t* = *t*_*n*_ − *t*_*n*–1_, and 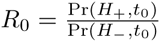.

### Appendix B: The continuum limit for the log likelihood ratio of two alternatives

In this section, we derive a continuum equation for the log likelihood ratio *y*_*n*_ := ln *R*_*n*_. We will proceed by first defining a family of stochastic difference equations for *y*_*n*_, which are parameterized by the timestep Δ*t* = *t*_*n*_ − *t*_*n*−1_, between pairs of observations. By choosing an appropriate parameterization, we obtain a continuum limit that is a SDE. To begin, we divide both sides of Eq. (A3) by *R*_*n*−1_ and take logarithms to yield

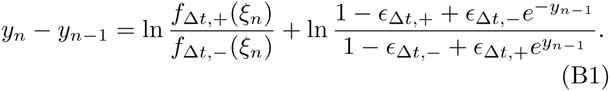

Following [2, 40], we assume that the time interval between individual observations, Δ*t,* is small. Denote by Δ*y*_*n*_ = *y*_*n*_ − *y*_*n*−1_ the change in the log likelihood ratio due to the observation at time *t*_*n*_. By assumption, the probability that the environment changes between two observations scales linearly with Δ*t* up to higher order terms, so that *∊*_Δ*t,±*_ := Δ*t∊_±_* + *o*(Δ*t*). Omitting higher order terms in Δ*t*, Eq. (B1) can then be rewritten as

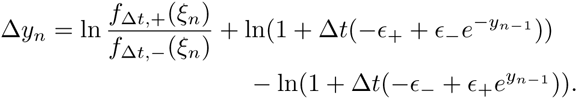

Since we assumed Δ*t* ≪ 1, we can use the approximation ln(1 +*a*) *≈ a* which is valid to linear order in |*a*| ≪ 1. We also assume that the change in the log likelihood ratio, Δ*y*_*n*_, is small over the time interval Δ*t*, so *y*_*n*−1_ can be replaced by *y*_*n*_ on the right-hand side of the equation. We obtain

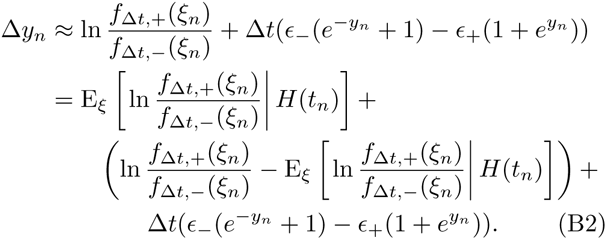

where we conditioned on the state of the environment, *H*(*t*_*n*_) = *H*_*±*_ at time *t*_*n*_. Replacing the index *n*, with the time *t,* we can therefore write

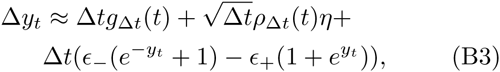

where *η* is random variable with standard normal distribution, and

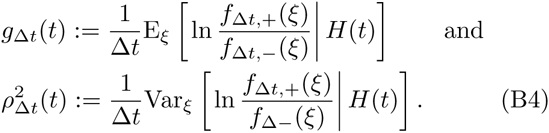

Clearly, the drift *g*_Δ*t*_ and variance 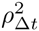 will diverge or vanish unless *f*_Δ*t,±*_(*ξ*) are scaled appropriately in the Δ*t →* 0 limit. We discuss different ways of introducing such a scaling in the next section.

Assuming that we have well-defined limits *g*(*t*) := lim_Δ*t→*0_ *g*_Δ*t*_(*t*) and 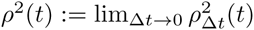, the discretetime stochastic process, Eq. (B3), approaches the SDE

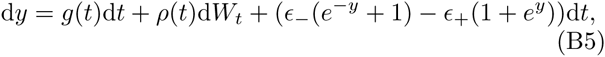

where *W*_*t*_ is a standard Wiener process. This limit holds in the sense of distributions. Roughly, the smaller Δ*t* is, the closer the distributions of the random variables *y*_*n*_ and *y*(*t*_*n*_) whose evolutions are described by Eq. (B1), and Eq. (B5), respectively. This correspondence can be made precise using the Donsker Invariance Principle [23].

In sum, Eq. (B5), can be viewed as an approximation of the logarithm of the likelihood ratio whose evolution is given exactly by Eq. (A3). For a fixed interval Δ*t*, the parameters of the two equations are related via Eq. (B4), and *ϵ*_Δ*t*,±_/Δ*t* = *ϵ*_±_.

### Appendix C: Precise correspondence

We now discuss two approaches in which the correspondence between Eqs. (B1) and (B5) can be made exact. We choose a specific scaling for the drift and variance arising from each observation, *ξ*_*n*_. Suppose that over the time interval Δ*t*, an observation, *ξ*_*n*_, is a result of *r*Δ*t* separate observations – for example the measurement of the direction of *r*Δ*t* different moving dots [3]. In this case the estimate of the average of the individual measurements – *e.g.* the average of the velocities of dots in a display – will have both a mean and a variance that increase linearly with Δ*t*.

As a concrete example we can compute *g*(*t*) and *ρ*(*t*) in SDE (B5) when observations, *ξ*_*n*_, follow normal distributions with mean and variance scaled by Δ*t*,

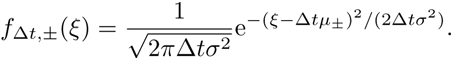

Using Eq. (B4) it is then straightforward to compute [2, 40],

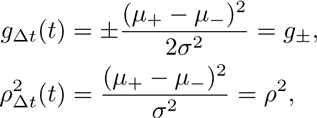

and note that *g*(*t*) ∈ {*g*_+_, *g*_−_} is a telegraph process [25] with the probability masses *P* (*g*_+_, *t*) and *P* (*g*_−_, *t*) evolving according to the master equation *P*_*t*_(*g*_*±*_, *t*) = ∓*E*_+_*P* (*g*_+_*, t*) *± E_−_P* (*g*_−_, *t*). In this case *ρ*^2^(*t*) = *ρ*^2^ remains constant.

More generally, we can obtain an identical result by considering that each observation made on a time interval consists of a number of sub-observations, each with statistics that scale with the length of the interval and the number of sub-observations. We define a family of stochastic processes parameterized by *k*, the number of sub-observations made in an interval of length Δ*t*. As above, when *k* = 1, we assume that an observation *ξ*_*n*_ is the result of *r*Δ*t* separate observations. Assuming *r* is large, note that for *k >* 1 each of the *k* subobservations contain roughly *r*_*k*_ = [*r*Δ*t/k*] observations with mean and variance that scale linearly with *r*_*k*_ ∝ Δ*t/k*. We can achieve this by approximating ln 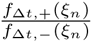 in Eq. (B2) by the family of stochastic processes parameterized by *k* [2]

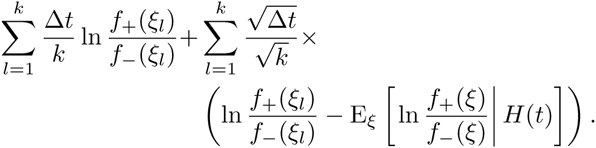

The scaling in this approximation guarantees that the drift is given by the limit 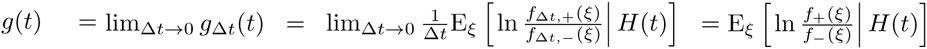 and the variance 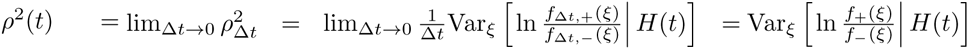 Furthermore, as *k* → ∞, by the Central Limit Theorem,

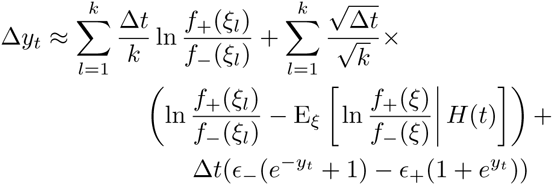

converges in distribution to

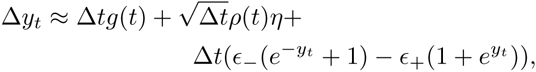

where *η* is a standard normal random variable. Taking the limit Δ*t* → 0 yields Eq. (B5).

### Appendix D: Continuum limit for log probabilities with multiple alternatives

We now describe the calculation of the continuum limit of the recursive system defining the evolution of the probabilities *L*_*n,i*_ = Pr(*H*(*t*_*n*_) = *H*_*i*_|*ξ*_1:*n*_) of one among multiple alternatives (environmental states), *H*_*i*_, *i* = 1,.., *N*. The state of the environment, and equivalently the correct choice at time *t*, again change stochastically. We assume that the transitions between the alternatives are memoryless, with transition rates ϵ_Δ*t,ij*_ := Pr(*H*(*t*_*n*_) = *H*_*i*_ | *H*(*t*_*n*−1_) = *H*_*j*_). Using Bayes’ rule and rearranging terms (analogous to the derivation of Eqs. (A1) and (A2)), we can express each probability *L*_*n,i*_ in terms the probability at the time of the previous observation, *L*_*n*−1*,j*_,

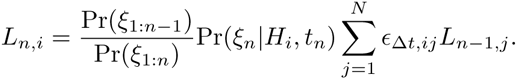

Since we are only interested in comparing the magnitude of the probabilities, we can drop the common prefactor 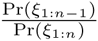, and use the fact that 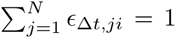 (since ϵ_Δ*t,ij*_ is a left stochastic matrix) to write ϵ_Δ*t,ii*_ = 1 − *Σ*_*j≠i*_ *ϵ*_Δ*t,ji*_ and obtain

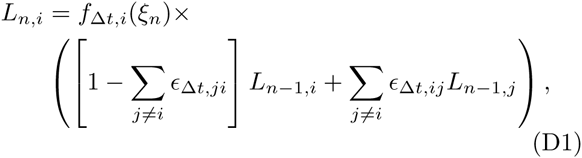

where *f*_Δ*t,i*_(*ξ*_*n*_) = Pr(*ξ*_*n*_*|H*_*i*_, *t*_*n*_). From Eq. (D1), it follows that log of the rescaled probabilities, *x*_*i*_ := ln *L*_*i*_, satisfies the recursive relation

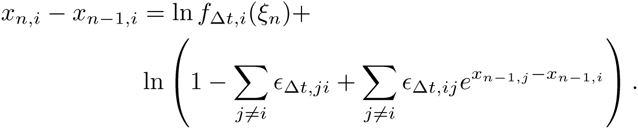

To derive an approximating SDE, we denote by Δ*x*_*n,i*_ = *x*_*n,i*_ − *x*_*n−*1,*i*_, the change in the log probability due to an observation at time *t*_*n*_. As before, we assume ϵ_Δ*t,ij*_ := Δ*t ϵ*_*ij*_ + *o*(Δ*t*) for *i* ≠ *j*, and drop the higher order terms, giving

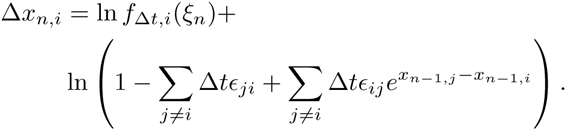

Assuming Δ*t* ≪ 1, we again use the approximation ln(1+ *a*) *≈ a* for *|a|* ≪ 1. We also assume that the change in the log probability, *|*Δ*x*_*n,i*_*|* ≪ 1, is small over the time interval Δ*t*, so that

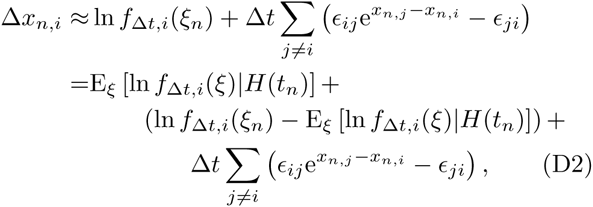

where we condition on the current state of the environment *H*(*t*_*n*_) ∈ {*H*_1_*,…, H*}.

Replacing the index *n*, by the time *t*, we can therefore write

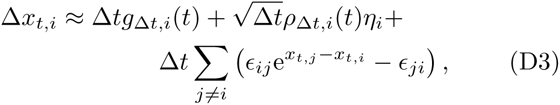

where *η*_*i*_’s are correlated random variables with standard normal distribution

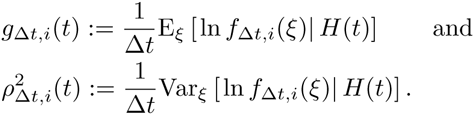

The correlation of *η*_*i*_’s is given by

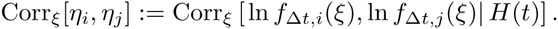

Note that Eq. (D3) is the multiple-alternative version of Eq. (B3). Equivalently, we can write Eq. (D3) as

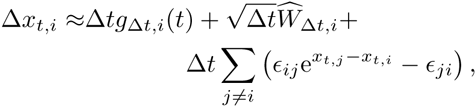

where 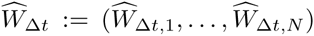 follows a multivariate Gaussian distribution with mean zero and covariance matrix Σ_Δ*t*_ given by

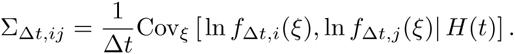

Finally, taking the limit Δ*t →* 0, and assuming that the limits

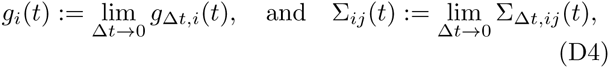

are well defined, we obtain the system of SDEs

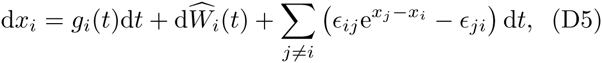

or equivalently as the vector system

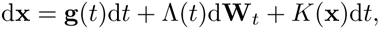

where **g**(*t*) = (*g*_1_(*t*),…, *g*_*N*_ (*t*))^*T*^ and Λ(*t*)Λ(*t*)^*T*^ = Σ(*t*) are defined using the limits in Eq. (D4), *K*_*i*_(**x**) = ∑*j*≠*i* (*ϵ*_*ij*_^e*x*_*j*_ −*x*_*i*_^ − *ϵ*_*ji*_), and the components of **W**_*t*_ are independent Wiener processes. We can recover Eq. (B5) by taking *N* = 2, letting *y* = *x*_1_ *− x*_2_, and exchanging the indices 1 and 2 with + and *, −* respectively.

As in the case of two alternatives, Eq. (D5) can be viewed as an approximation of the logarithm of the probability whose evolution is given exactly by Eq. (D1). For a fixed interval Δ*t*, the parameters of these equations are related via Eq. (D5), and ϵ_Δ*t,ij*_*/*Δ*t* = *ϵ*_*ij*_.

The limits *g*_*i*_(*t*) := lim_Δ*t→*0_ *g*_Δ*t,i*_(*t*) and Σ_*ij*_(*t*) := lim_Δ*t→*0_ Σ_Δ*t,ij*_(*t*) are defined when the statistics of the observations scale with Δ*t*. As we argued above, this can be obtained by considering observations drawn from a normal distribution with mean and variance scaled by Δ*t*:

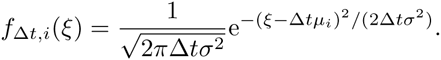

Alternatively, the required scaling can also be obtained when each observation made on a time interval consists of a number of sub-observations, (*ξ*_1_,…, *ξ*_*k*_), with mean and variance scaled by 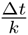. To do so we approximate ln *f*_Δ*t,i*_(*ξ*_*n*_) in Eq. (D2) by

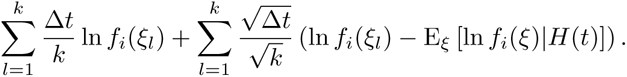

### Appendix E: Log likelihood ratio for multiple alternatives

We can also derive a continuum limit for the log likelihood ratio for any two choices *i, j ∈ {*1, 2*,…, N}*. From Eq. (D1), the likelihood ratio *R*_*n,ij*_ = *L*_*n,i*_/*L*_*n,j*_. We note that this will provide us with a matrix of stochastic processes. We start with the recursive equation

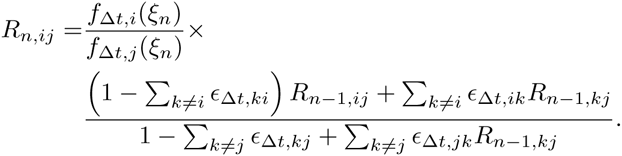

We can thus derive the continuum equation for the log likelihood ratio *y*_*n,ij*_ := ln *R*_*n,ij*_, as we did in the case of two alternatives. Since *y*_*ij*_(*t*) is the difference *y*_*ij*_(*t*) = *x*_*i*_(*t*) − *x*_*j*_(*t*), from Eq. (D5) we obtain

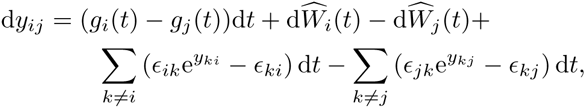

Or

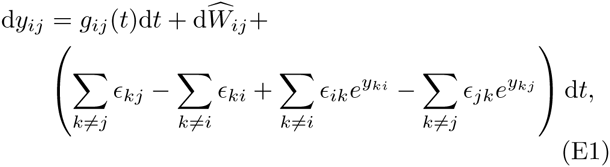

where 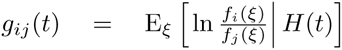 and 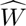 is a Wiener process with covariance matrix given by 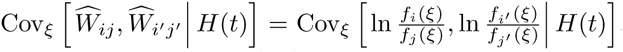. We can also write Eq. (E1) in vector form

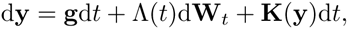

where 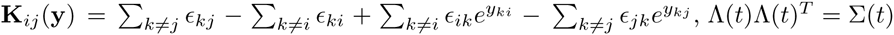 is the covariance matrix, and the components of **W**_*t*_ are independent Wiener processes.

### Appendix F: Log probabilities for a continuum of alternatives

Finally, we examine the case where an observer must choose between a continuum of hypotheses *H*_*θ*_ Where *θ* ∈ [*a, b*]. Thus, we will first derive a discrete recursive equation for the evolution of the probabilities *L*_*n,θ*_ = Pr(*H*(*t*_*n*_) = *H*_*θ*_*|ξ*_1:*n*_). The state of the environment, the correct choice at time *t*, again changes according to a continuous time Markov process. We define this stochastically switching process through its transition rate function ϵ _Δ*t,θθ*_ *′*, which is given for *θ′ ≠ θ* as

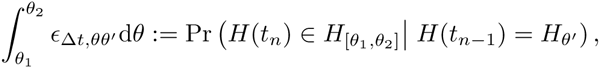

where *H*_[*θ*_1_*,θ*_2_]_ is the set of all states *H*_*θ*_ with *θ* in the interval [*θ*_1_*, θ*_2_]. Thus, ϵ _Δ*t,θθ*_*′* describes the probability of a transition over a timestep, Δ*t*, from state *H*_*θ*_ *′* to some state *H*_*θ*_, with *θ* ∈ [*θ*_1_*, θ*_2_]. This means that 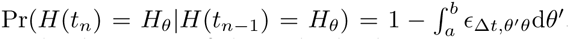 As in the derivation of the multiple alternative 2 *≤ N < ∞* case, we can express each probability *L*_*n,θ*_ at time *t*_*n*_ in terms of the probabilities *L*_*n−*1*,θ*_′ at time *t*_*n−*1_, so

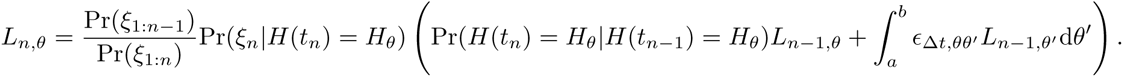

Notice that the sum from the *N* < ∞ case, as in Eq. (D1), has been replaced with an integral over all possible hypotheses *H*_*θ*_*′*, *θ ′* ∈ [*a, b*] and a term corresponding to the probability of the environment not changing. Again we drop the common factor 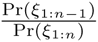, since we wish to compare the magnitudes of the probabilities. We obtain

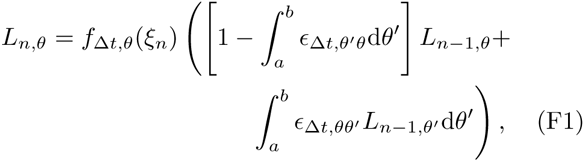

where *f*_Δ*t,θ*_(*ξ*_*n*_) = Pr(*ξ*_*n*_ | *H*(*t*_*n*_) = *H*_*θ*_). From Eq. (F1), we can thus derive a recursive relation for the log of the rescaled probabilities *x*_*n,θ*_ := ln *L*_*n,θ*_ in terms of *x*_*n−*1*,θ*_ so

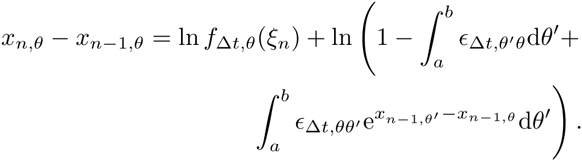

To approximate this discrete-time stochastic process with a SDE, we denote by Δ*x*_*n,θ*_ = *x*_*n*,θ_ – *x*_*n−*1,*θ*_, the change in log probability due to the observation at time *t*_*n*_. Furthermore, we assume ϵ _Δ*t,θθ*_*′* := Δ*tE*_*θθ*_′ + *o*(Δ*t*) and drop higher order terms,

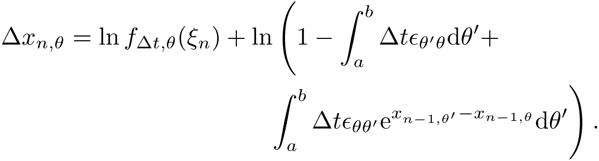

Assuming Δ*t* ≪ 1, we again utilize the approximation ln(1 + *a*) *≈ a* for *|a|* ≪ 1. Assuming *|*Δ*x*_*n,θ*_*|* ≪ 1,

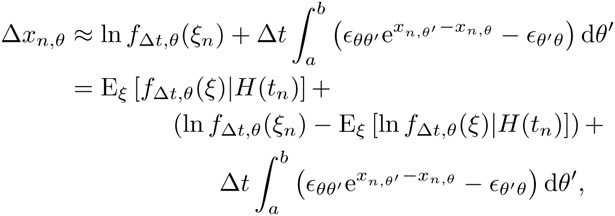

conditioned on the current state of the environment *H*(*t*_*n*_) = *H*_*φ*_ where *φ* ∈ [*a, b*].

Exchanging the index *n* with the time, *t*, we can therefore write

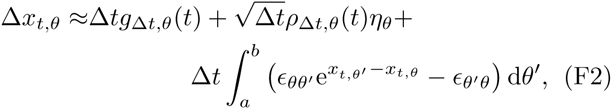

where *η_θ_*’s are correlated random variables which marginally follow a standard normal distribution, and

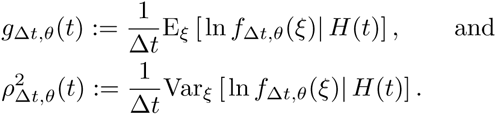

The correlation of *η*_*i*_’s is given by

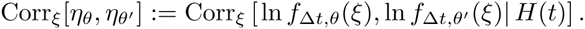

Equivalently, we can write Eq. (F2) as

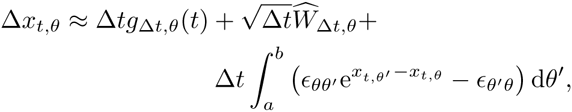

where 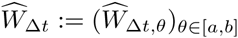 follows a multivariate Gaussian distribution with mean zero and covariance function Σ_Δ*t,θθ*_ *′* given by

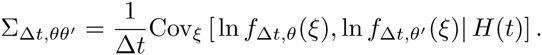

Finally, taking the limit Δ*t* → 0, and assuming that the limits

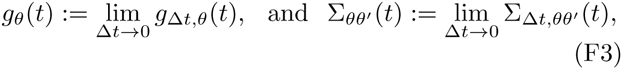

are well defined, we obtain the system of SDEs

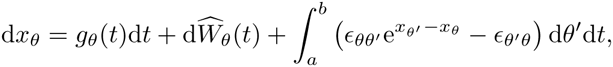

or equivalently as the system of SDEs

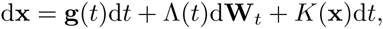

where **g**(*t*) = (*g*_*θ*_(*t*))_*θ*∈[*a,b*]_ and Λ(*t*)Λ(*t*)^*T*^ = Σ(*t*) are defined using the limits in Eq. (F3), 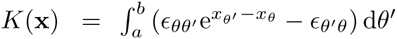, and the components of **W**_*t*_ are independent Wiener processes.

